# Spectral fingerprints or spectral tilt? Evidence for distinct oscillatory signatures of memory formation

**DOI:** 10.1101/373514

**Authors:** Marie-Christin Fellner, Stephanie Gollwitzer, Stefan Rampp, Gernot Kreiselmeyr, Daniel Bush, Beate Diehl, Nikolai Axmacher, Hajo Hamer, Simon Hanslmayr

## Abstract

Decreases in low frequency power (2-30 Hz) alongside high frequency power increases (>40 Hz) have been demonstrated to predict successful memory formation. Parsimoniously this change in the frequency spectrum can be explained by one factor, a change in the tilt of the power spectrum (from steep to flat) indicating engaged brain regions. A competing view is that the change in the power spectrum contains several distinct brain oscillatory fingerprints, each serving different computations. Here, we contrast these two theories in a parallel MEG-intracranial EEG study where healthy participants and epilepsy patients, respectively, studied either familiar verbal material, or unfamiliar faces. We investigated whether modulations in specific frequency bands can be dissociated in time, space and by experimental manipulation. Both, MEG and iEEG data, show that decreases in alpha/beta power specifically predicted the encoding of words, but not faces, whereas increases in gamma power and decreases in theta power predicted memory formation irrespective of material. Critically, these different oscillatory signatures of memory encoding were evident in different brain regions. Moreover, high frequency gamma power increases occurred significantly earlier compared to low frequency theta power decreases. These results speak against a “spectral tilt” and demonstrate that brain oscillations in different frequency bands serve different functions for memory encoding.

## Introduction

Understanding the neural processes that mediate encoding of new memories is fundamental. Encoding processes are at the first stage of transforming transient experiences into memories, which essentially make us who we are. The subsequent memory paradigm allows investigating these processes by contrasting neural activity during later remembered events with activity during later forgotten events at encoding [1]. Electrophysiological methods, like intracranially recorded EEG or EEG/MEG, are particularly promising to offer a mechanistic understanding of memory formation processes. M/EEG and iEEG index neural synchronization and desynchronization processes [2,3], which have been directly linked to synaptic plasticity [4–6]. A number of subsequent memory studies demonstrate that memory formation is indicated not by one particular frequency, but instead by complex changes in multiple frequencies ranging from 2-100 Hz, encompassing theta, alpha, beta and gamma frequency bands [7,8]. A common finding is that power decreases in low frequencies (<30 Hz) paired with increases in high frequency power (>40 Hz) are beneficial for memory formation. This change in the spectral pattern can result from two different processes: (i) a change in the “spectral tilt”, i.e. a shift from low frequency activity to high frequency activity [9–11] or (ii) changes at multiple distinct frequency bands related to distinct subprocesses involved in memory formation. Here we contrast these two frameworks in a subsequent memory paradigm and show that memory-related frequency components can be dissociated on three levels: experimental condition, temporal dynamics and brain regions.

Decreases in low frequency power are often accompanied by increases in high frequency power during various tasks. This is especially true for alpha/beta power decreases and gamma power increases [12–14]. Human EEG shows a 1/f-like characteristic whereby power decreases with increasing frequency [11,15]. A change in the “tilt” of this 1/f spectrum can parsimoniously explain such low frequency decreases accompanied by gamma power increases [8,11,16]. The “spectral tilt” idea makes no assumptions about synchronization processes, but instead views a shift from low to high frequency activity as a proxy for increased neural firing [10,17,18]. Evidence for the spectral tilt theory comes from recent brain stimulation studies which were able to boost memory encoding by stimulating specifically during periods characterized by “bad memory states”, i.e. increased low and decreased high frequency activity[19,20].

The spectral tilt framework makes three specific predictions, which will be tested here: (i) increases in gamma power should be accompanied by decreases in alpha/beta and theta power and vice versa. Therefore, it should not be possible to experimentally dissociate gamma power increases from alpha/beta and theta power decreases; (ii) decreases in low frequency (alpha/beta/theta) power and increases in high frequency (gamma) power should occur in strongly overlapping brain regions; and (iii) low frequency decreases and high frequency increases should occur at the same time. In sum the “tilt” assumption suggests that high frequency increases and low frequency decreases reflect the same process. Following this assumption many iEEG studies confine analysis to broadband high frequency power changes, presuming that high frequency increases reliably map brain activity [21–25]

The “spectral fingerprints” framework assumes that oscillatory changes in different frequencies indicate different neural processes in different brain regions [26], each reflecting a specific function in the service of memory [7,27,28]. The framework of “spectral fingerprints” has a specific relevance when studying memory formation, because the idea that several distinct subprocesses contribute to memory formation is integral to many memory models [29–31]. Prior work has linked different oscillatory changes to assumed memory subprocesses. For instance, theta oscillations have been related to binding processes in a medial temporal Lobe (MTL) network [32–34], whereas alpha/beta power decreases have been related to semantic processing during memory encoding [35–37]. Gamma power increases in sensory areas have been suggested to reflect locally synchronized activity and to indicate bottom-up sensory processing [3,38]. The “spectral fingerprints” framework therefore arrives at very different predictions compared to the spectral “frequency tilt” framework. Specifically, it suggests that it should be possible to experimentally dissociate gamma power increases from theta and alpha/beta power decreases. It further suggests that the power changes of different frequency bands could occur in different brain regions and at different time points.

To test the two frameworks against each other, we employed the same memory task while recording iEEG in patients and MEG in healthy participants. It is important to note that many subsequent memory studies arguing in favor of spectral fingerprints used non-invasive MEG/EEG [32,37] whereas most studies finding evidence for a spectral tilt used intracranially recorded EEG [8,20]. To investigate whether specific spectral fingerprints can be dissociated by varying encoding demands, we utilized the well-established finding that memory encoding of words is crucially different from encoding of unfamiliar faces. Specifically encoding of words heavily depends on semantic processing [39], whereas encoding of unfamiliar faces solely depends on visual processing [40,41]. Therefore, we recorded MEG and iEEG during encoding of words and faces (see Figure 1A) to rule out differences in recording methods as a confounding factor (i.e. higher sensitivity of iEEG to capture high frequency dynamics [42,43]. This combined measurement also allowed us to capture changes in a large frequency range (2-100 Hz) on a whole brain and at a more local level.

**Figure 1:**
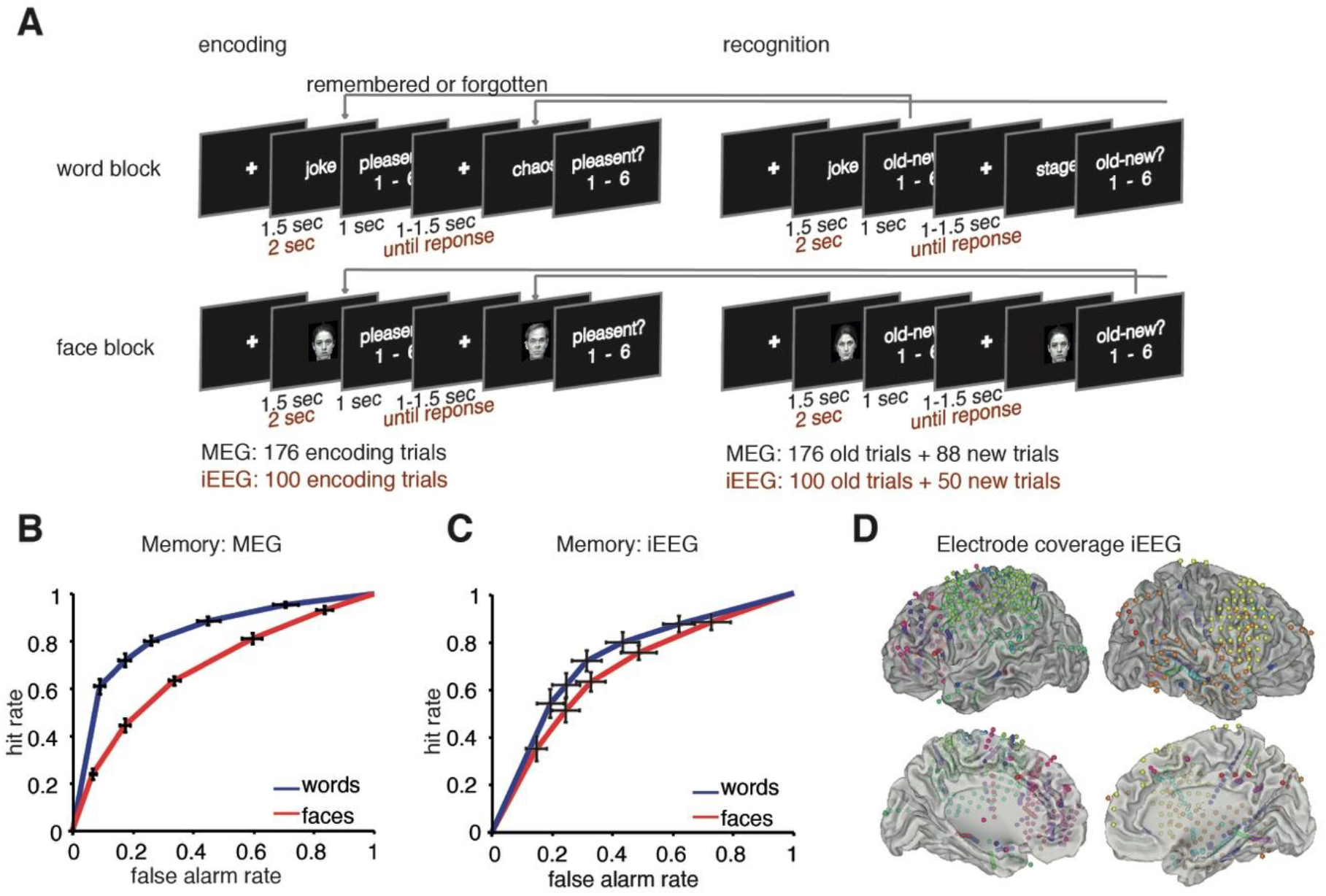
(A) Paradigm. The experiment was split in two blocks: one block word encoding and recognition and one for faces respectively. The paradigm was slightly adapted for the iEEG patient sample. Behavioral performance in the MEG dataset (B) and iEEG dataset (C). ROC curves show memory performance for faces and words, error bars plot SEM for each rating. A left upward shift of the ROC indicates higher recognition performance, i.e. more hits and less false alarms. (D) Electrode coverage in the iEEG patient sample: The bipolar referenced virtual electrode locations of each contact included in the reported analysis are plotted here, colors code different patients.

## Results

### Behavioral results

Memory performance is shown by means of ROC curves in Figures 1B and 1C, depicting memory performance in the MEG sample (healthy participants) and the iEEG sample (patients). Recognition performance in the MEG sample was higher for words (mean d’=1.99) than faces (mean d’=0.83, t(19)=7.836 p<.0001). A similar effect was observed in the iEEG sample which just fell short of significance likely due to the higher variance in performance between patients (mean d’ words=1.37 vs. mean d’ faces=0.90; t(12)=2.12 p=0.052). These results are in line with previous studies demonstrating the difficulty of memorizing unfamiliar faces [41,44]. The difference in recognition performance already hints at different processes involved in encoding of words in contrast to faces.

### Material specific effects: lower frequency bands

The experiment involved two main factors: material (words vs. faces) and subsequent memory (remembered vs. forgotten). In a first step we analyzed overall differences of material, i.e. differences in power between processing a word or face item independent of memory. This allowed us to identify time-frequency windows of interest for later analysis, and served as a first test to check for co-localized low and high frequency power decreases/increases.

Sensor level analysis of the MEG data revealed significant differences in the lower frequencies, i.e. the alpha/beta band. A cluster permutation statistic across sensors, frequency-bands (2-30Hz) and time (0 to 1.5 sec post stimulus) revealed two significant clusters spanning alpha/beta frequency bands (word<face, p_corr_=.007, word>face, p_corr_=.002 see Figure 2A): One cluster exhibited relative greater alpha/beta power decreases (8-20Hz) for words than faces, located at left frontal sensor sites at 0.3-1.5 sec poststimulus. A second cluster exhibited relatively larger alpha/beta power decreases (8-20Hz) for faces than words and was located at posterior sensors in the same time interval (0.3-1.5 sec). In accordance with the sensor level results, source analysis revealed significant clusters in regions commonly involved in word and face processing, respectively (Figure 2B): Stronger alpha/beta power decreases for words compared to faces (shown in red) were localized to areas typically involved in semantic processing encompassing the inferior and middle frontal gyrus, supramarginal gyrus, Heschl’s gyrus and temporal pole and middle and superior temporal gyrus (peak t(19) =-4.16 at MNI -54, 0, 50 corresponding to left precentral gyrus, word<face p_corr_=0.049, t-sum=-826.60). Stronger alpha/beta power decreases for faces compared to words (shown in blue) were localized to areas typically involved in visual processing (Figure 2B), spanning lingual, occipital middle, fusiform, temporal middle and inferior gyrus in the right hemisphere. The peak of the source localization was in right inferior occipital gyrus (t(19)=9.04, MNI 46, 80, 10, source cluster: p_corr_ <0.001, t-sum=62700.7), In summary, the material effects in MEG are in line with the hypothesis that areas involved in material-specific processing of stimuli show a relative decrease in alpha/beta power. Decreases in alpha/beta power seem to closely map to areas involved in face and word processing as found with fMRI [45]

**Figure 2:**
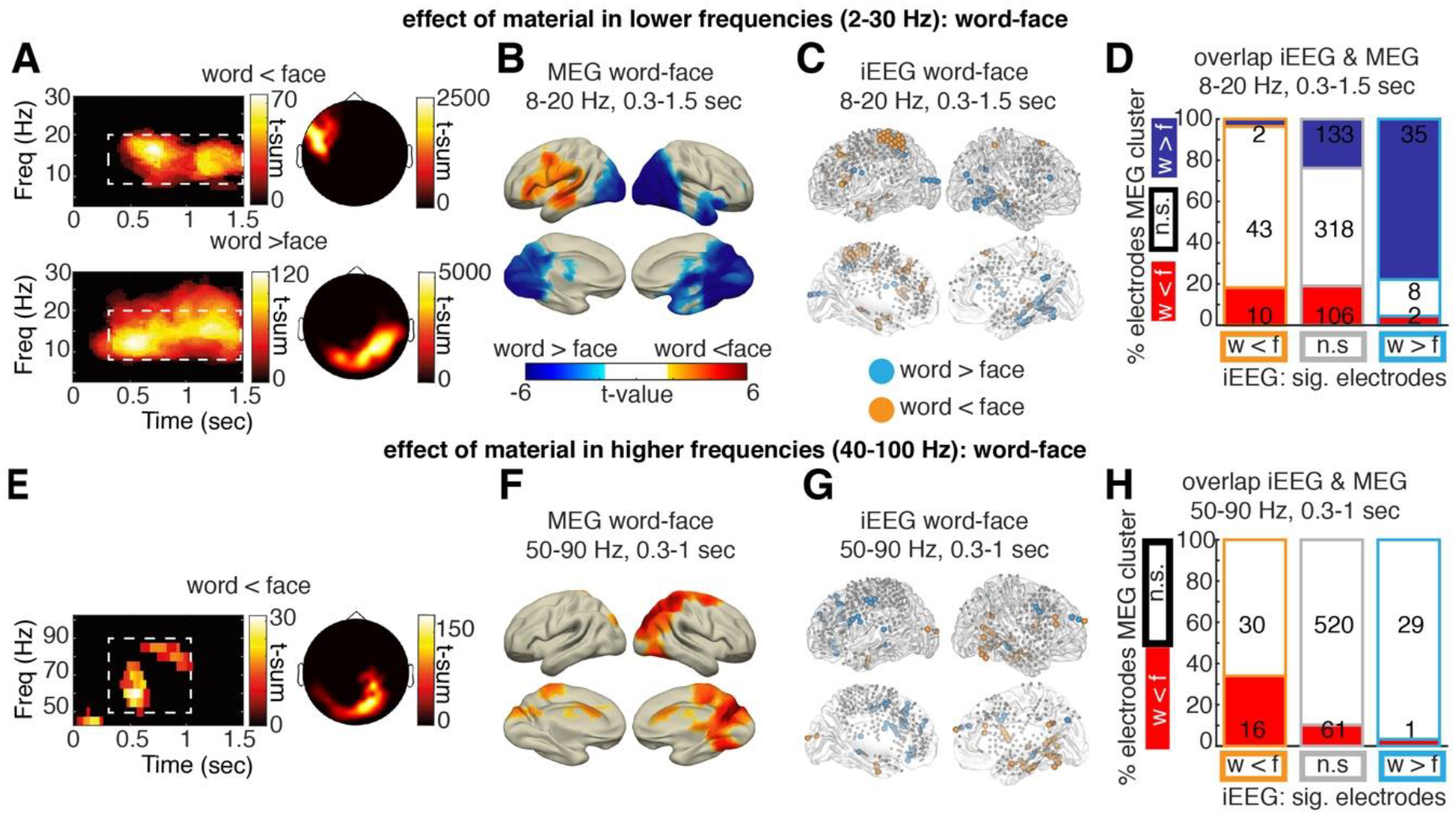
MEG Main effect of material: word-faces. (A) Main effect of material: word vs face condition irrespective of memory: Significant clusters (p_corr_<0.05) returned by a cluster permutation statistic clustering across sensors, frequencies and channels. One cluster showed a greater decrease in alpha/beta power for words relative to faces at left frontal sensors, another cluster showed a stronger decrease in alpha/beta power for faces relative to words at posterior sensors. (B) Significant sources (p_corr_<0.05) showing a main effect of material in alpha/beta power in the time-frequency window identified in sensor level data. (C) Significant electrodes (p<0.05, uncorrected) in iEEG showing a significant change in alpha/beta power in the respective time-frequency window identified in MEG. (D) The overlap of MEG and iEEG material effects is visualized by plotting for each category of iEEG electrodes (w<f: light blue, w>f: orange, or no sig difference: grey), the percentage of electrodes located in areas of significant MEG source clusters (clusters of w<f: red, w>f: blue, or no significant difference: white). The numbers plotted on top of the bars denote the absolute numbers of electrodes in each source area respectively. (E-H) show the corresponding analysis in the high frequency range. No significant MEG cluster emerged for word > face.

In a next step we assessed whether similar material specific effects occurred in the iEEG data. To this end, the main effect of material in the time-frequency window identified in the MEG analysis (i.e. 8-20 Hz, 0.3-1.5 sec) was calculated in each single electrode across the whole patient sample. Figure 2C shows all electrodes exhibiting a significant main effect of material (p<0.05, uncorrected) in the alpha/beta frequency range, as assessed by a separate ANOVA with the factors material and memory in each electrode. Electrodes showing stronger alpha/beta power decreases for words relative to faces were spread across the left hemisphere (highlighted in orange in Figure 2C). In contrast, electrodes showing stronger alpha/beta power decreases for faces relative to words are predominantly located in the right posterior and ventral visual stream (highlighted in light blue in Figure 2C). The location of MEG sources and the distribution of significant iEEG electrodes for the words vs faces contrast showed a good degree of overlap visually (Figure 2 A-C). To formally test this overlap, a χ^2^ test of independence was calculated by counting the number of significant iEEG electrodes (Figure 2C) separately for areas inside and outside of significant source clusters in MEG source analysis (Figure 2B). The localization of significant iEEG electrodes and the MEG source clusters was not independent (Figure 2D, χ^2^(4) =79.79, p<0.0001): iEEG electrodes were more likely to exhibit significant modulation of alpha/beta power (marked in light blue and orange) if located in regions that exhibit the same modulation in MEG (marked in blue and red). This finding demonstrates an overlap of iEEG and MEG results, despite the different properties of iEEG and MEG in spatial resolution and spatial sampling.

### Material specific effects: higher frequency bands

Material specific effects in the higher frequency range (40-100Hz) were analyzed congruently to effects in the lower frequency range. First, time-frequency windows of interest were identified via an open cluster permutation statistic across all MEG sensors, in a frequency range from 40-100Hz and a time range from 0-1.5 sec poststimulus. This analysis revealed three clusters exhibiting a significant increase in gamma power related to faces in contrast to words, all located at posterior sensors (~50-90 Hz, 0.3-1.0 sec post stimulus, p_corr_=.011, p_corr_=.011, p_corr_= .046, Figure 2E). These face-driven gamma power increases were localized to areas involved in visual processing (right cuneus, occipital superior gyrus, superior parietal gyrus and precuneus, peak in right cuneus, t(19)=-8.07, MNI 16, -80, 30, source cluster p<0.001, t-sum=-2797.9). This result is consistent with the hypothesis that task active regions can be identified by increases in high frequency power. However, this result was specific to face stimuli, as no clusters showing significant high frequency power increases for word stimuli were identified.

Analogous to the low frequency power analysis, the main effect of material in iEEG data was analyzed in the time-frequency window identified in the MEG data (50-90 Hz, 0.3-1 sec). In Figure 2G all electrodes showing significant increases in high frequency (gamma) power for faces compared to words are highlighted in orange, and all electrodes showing gamma power increases for words are shown in blue (all p<0.05, uncorrected). To quantify the overlap between the two modalities a χ^2^ test of independence was calculated. Here, only electrodes with power increases for faces compared to words were tested, since no gamma power increases for words were found in the MEG data. The number of iEEG electrodes showing significant relative power increases for faces was higher in areas exhibiting the same effect in MEG (Figure 2H, χ^2^(2) =26.25, p<0.0001). Thus, changes in gamma power show a similar concordance of MEG and iEEG data as changes in alpha/beta power. Interestingly, in occipital areas decreases in alpha/beta power seem to co-occur with increases in gamma power, while no such relationship was evident for areas related to the processing of words.

### Material-specific high frequency effects in low frequency clusters

Material specific effects were found in the form of lower frequency power decreases in the alpha/beta range and high frequency gamma power increases. In line with the spectral tilt hypothesis, our results suggest that gamma power increases are related to decreases in alpha/beta power in occipital areas. However, no comparable increase in gamma power was evident in areas involved in word processing in the MEG using whole brain statistics. To assure that such word-related high frequency (gamma) power increases were not overlooked because of a lack of statistical power or weaker signal-to-noise ratio, we employed a region of interest analysis focusing on gamma changes in MEG sources and iEEG electrodes exhibiting material related alpha/beta power changes (Figure 3). High frequency effects of material were assessed exclusively in MEG source clusters exhibiting material related alpha/beta power changes (see Figure 2B for the respective clusters). In iEEG, an examination of gamma power changes was restricted to electrodes exhibiting material related alpha/beta power changes. To calculate a random effects group statistic in the iEEG data (that is comparable to the MEG data), power spectra were averaged in each patient across all electrodes exhibiting a significant negative or positive main effect of material in alpha/beta power. For both modalities we then compared gamma power effects in those clusters (MEG) or electrodes (iEEG) exhibiting material specific low frequency power decreases.

**Figure 3:**
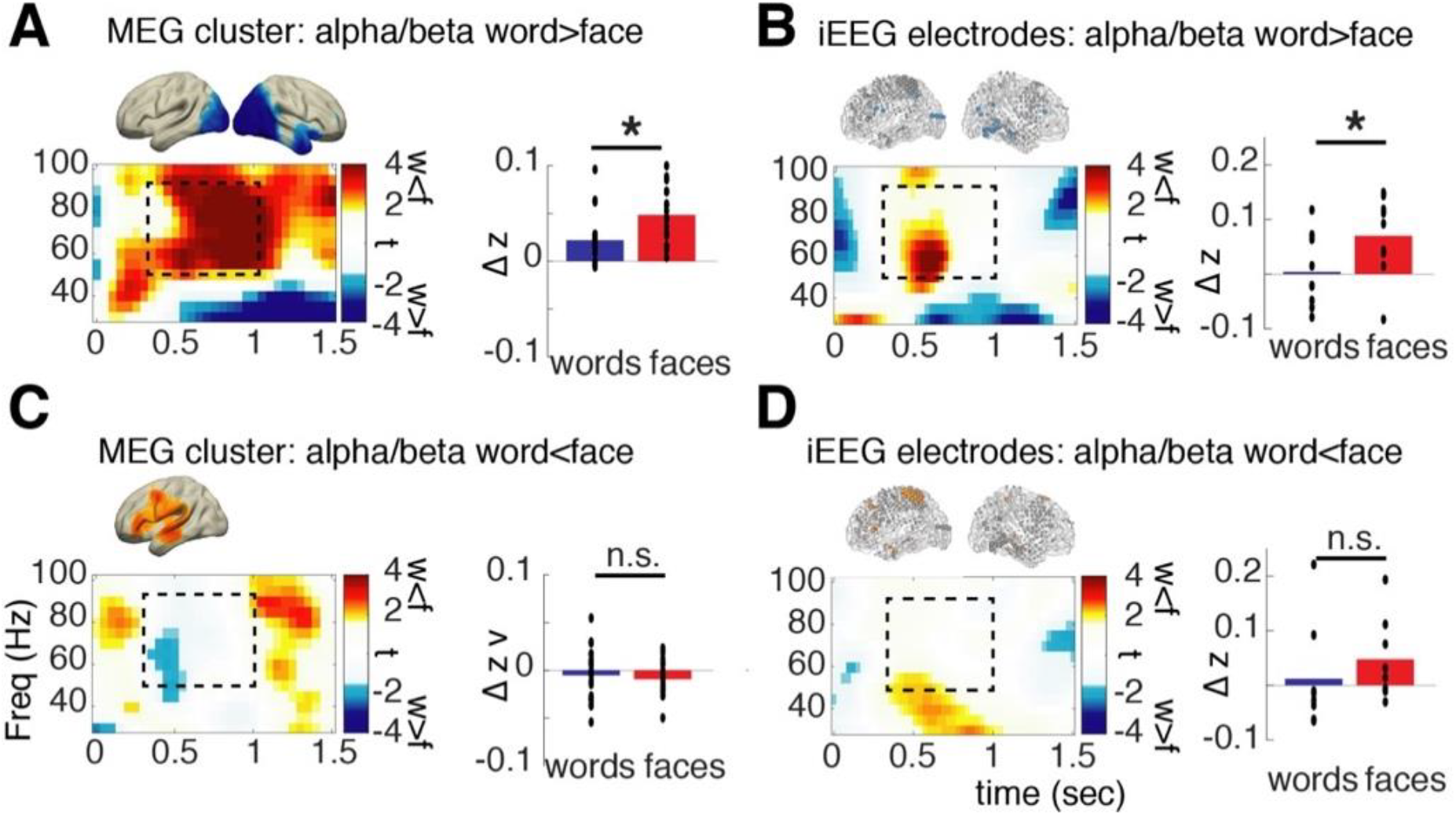
Material specific high frequency power changes in MEG source clusters and iEEG electrodes exhibiting material specific power changes in the alpha/beta range. Time frequency plots show t-values for the contrast of words vs. faces in MEG source clusters (A, C) and in iEEG electrodes (B, D). Bar plots show average power values for the word and face condition separately for the time-frequency window marked with dashed boxes, black dots show the respective single subject averages. Asterisks mark significant differences depending on material (p<0.05). Brain plots are replicated from Figure 2 to highlight the respective ROIs.

In the clusters/electrodes which exhibited face specific alpha/beta power decreases a concurrent increase in gamma power was evident, which parallels the above findings. Specifically, both modalities, MEG and iEEG, showed a significantly stronger gamma power increase for faces compared to words (Figure 3A & 3B, MEG: t(19)=-5.733, p<0.0001, iEEG: t(8)=-2.337, p=0.0476). Concerning the clusters/electrodes exhibiting word specific alpha/beta power decreases, however, no co-localized increases in high frequency (gamma) power were evident (Figure 3C & 3D, MEG: t(19)=0.60, p=0.56, iEEG: t(7)=-1.260, p=0.283). This pattern of results shows that the co-occurrence of gamma increases, and alpha/beta power decreases may be limited to specific brain areas. In posterior sensory processing regions, our results agree with the spectral tilt hypothesis, showing increases in gamma power co-occurring with decreases in low frequency power. However, in left lateralized areas exhibiting strong alpha/beta power decreases specific to word processing, no concurrent increases in gamma power were evident, neither in MEG nor in the more spatially resolved iEEG data, thus violating a key prediction of the spectral tilt hypothesis.

### Subsequent memory effects

Subsequent memory effects (SMEs, i.e. effects of memory encoding) were investigated in two time-frequency windows obtained in the above described material specific contrasts and described above, which are orthogonal to the memory contrast: the time-frequency windows showing alpha/beta (8-20 Hz, 0.3-1.5 sec) and gamma power changes (50-90 Hz, 0.3-1.0 sec). Additionally, theta power changes were investigated in a third time-frequency window from 2-5 Hz and from 1.0-1.5 sec based on prior studies [33,46]. We were specifically interested in interactions of memory and material, i.e. whether SMEs in specific frequency bands vary with material.

Since MEG and iEEG data crucially diverge in the brain coverage in each subject, different statistical analyses were employed. To parallelize analysis of iEEG and MEG data a two-stage procedure was used. In a first stage, regions of interest or electrodes of interest, which showed a main effect of memory (i.e. SMEs independent of material) were identified. In the second stage the data within these ROIs were tested for Material x Memory interactions. Notably, this procedure does not artificially inflate statistical power for finding interactions – it slightly biases the results against finding interactions since only ROIs with a tendency for showing SMEs in the same direction across materials are considered. This procedure can readily be applied to MEG as well as to iEEG data, which is not trivial since locations of electrodes vary from patient to patient, whereas MEG data provides whole brain coverage in every subject. This analysis allows for random effects analysis of memory x material interaction of in iEEG data at the expense of limiting analysis to electrodes and patients exhibiting significant main effects of memory. Since this selection may theoretically distort the findings, an additional fixed effects analysis was run including all electrodes combined across all patients (657 electrodes, Figure 4G). The fixed effects analysis (FEA) tested whether the number of significant electrodes across all patients exceeded the number of significant electrodes in a randomly permuted sample.

**Figure 4:**
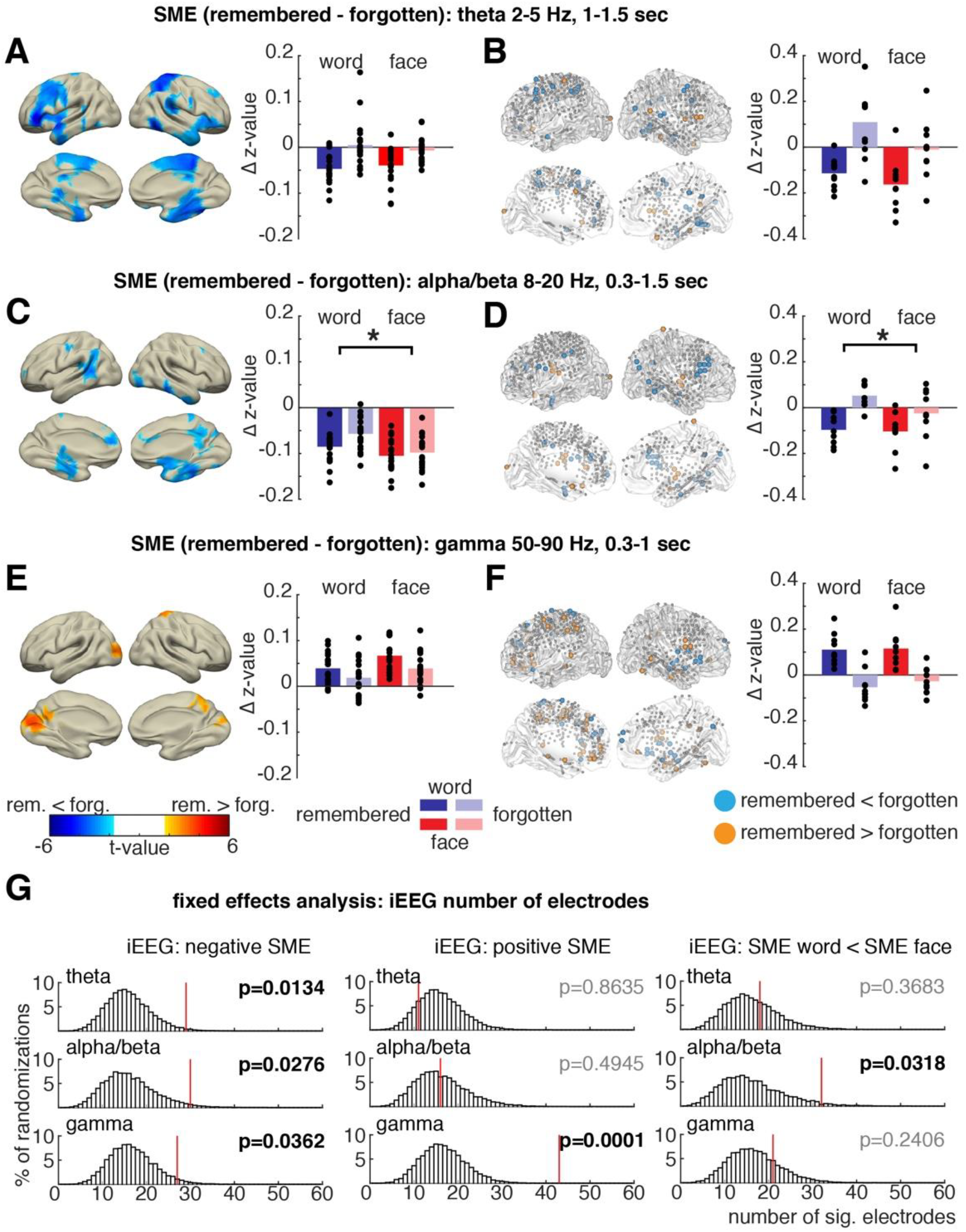
Material-independent and material-dependent subsequent memory effects. Subsequent Memory Effects (SME, remembered-forgotten) were analyzed in three time-frequency windows of interest. (A, C, E) show significant source clusters (p_corr_<0.05) for main effects of memory (SMEs irrespective of material). (B,D,F) iEEG electrodes exhibiting significant (p<0.05, uncorrected) positive or negative main effects of memory are highlighted in orange or blue respectively. Bar-plots on the right show average normalized power for all condition averages for the respective MEG source cluster or iEEG electrodes plotted to the left, respectively. Black dots show the respective single subject averages. Asterisks mark significant interaction effects (p<0.05). (G) Results of a fixed effects permutation analysis including all iEEG electrodes across all patients. The number of significant electrodes in the data exhibiting a significant effect is highlighted in red relative to the distribution of significant electrodes in randomly shuffled data.

### Theta

In MEG data decreases in theta power (2-5 Hz, 1-1.5 sec) were observed for later remembered compared to later forgotten items (source p<0.001, T-sum =-3446.8; Figure 4A). Theta power decreases spanned a temporo-cortical network including inferior frontal regions, parietal and temporal regions with a peak in the left inferior frontal gyrus (t(19)=-5.82, MNI -56, 40, -10). This negative theta SME did not differ between words and faces (interaction memory x material: t(19) =-1.02, p=0.32).

These effects were paralleled in the iEEG data: significant decreases in theta power were observed for later remembered compared to later forgotten items (i.e. negative SME) in the same time-frequency window (2-5 Hz, 1-1.5 sec, FEA: 29 electrodes, p_corr_ =0.0134, Figure 4G). No significant positive SME, i.e. increase in power for remembered vs forgotten, was found (FEA: 11 electrodes, p_corr_ =0.66). In electrodes exhibiting a significant negative theta SME no difference in SMEs depending on material across patients was evident (Figure 3B; analysis including 10 patients, 1-6 electrodes per patient, p=0.13, t(9) =-1.66). In the fixed effects analysis across all electrodes this result was replicated, no significant material dependent difference of SMEs was found (FEA: SME words < SME faces: p=0.37, 18 electrodes Figure 4G, SME words > SME faces: p_corr_ =0.75, 12 electrodes). Together, we observed the same pattern of results in MEG and iEEG. The data indicate that decreases in theta power predict memory formation independent of material.

### Alpha/beta

In the MEG data, decreases in alpha/beta power (8-20 Hz, 0.3-1.5 sec) predicted later memory (source level p= 0.005, t-sum = -2092.4). This negative SME was localized in areas encompassing typical encoding relevant regions: left superior frontal, inferior-medial temporal areas including the left hippocampus (peak right inferior temporal, t(19)=-4.75, MNI: 74, -40, -20; Figure 4C). Importantly, the alpha/beta power SMEs significantly differed between faces and words, as indicated by a significant Material by Memory interaction (p=0.020, t(19) =-4.38). Alpha/beta SMEs were significantly stronger for words in contrast to faces.

The same pattern of effects was evident in the iEEG data. Significant decreases in alpha/beta power were related to successful encoding across electrodes (FEA: alpha/beta: p_corr_ =0.028, 30 electrodes, Figure 4G), no significant increase in alpha/beta power during memory formation was found (FEA: p_corr_ =0.49, 16 electrodes, Figure 4G). Across patients with significant electrodes showing a significant negative alpha/beta SME, a similar significant difference in SMEs was evident as in MEG: here again a significant Material by Memory interaction was observed (p=0.0027, t(8) =-4.28, 9 patients, 1-7 electrodes per patient), which was driven by stronger power decreases for later remembered words compared to later forgotten words (Figure 4D). The same significant difference in SMEs between words and faces was obtained for alpha/beta power decreases across all electrodes (FEA: words<faces: p_corr_ = 0.032, 32 electrodes, SME words > SME faces: p_corr_ =0.66, 13 electrodes, Figure 4G).

These stronger negative alpha/beta SMEs for words than for faces in MEG and iEEG data, suggest a specific role of alpha/beta power decreases during memory formation for verbal material. This result further demonstrates that decreases in low frequency power do not uniformly contribute to later memory, since this interaction effect was specific to the alpha/beta band, but not evident in the theta range.

### Gamma

In the gamma time-frequency window (50-90 Hz, 0.3-1 sec) a significant positive SME, i.e. increases for later remembered items, emerged in the MEG data localized to occipital, and parietal regions (Figure 4E, p=0.048, t-sum=479.06). The peak of the source was located in left superior occipital gyrus (max t(19)=4.34, MNI -14, -90, 10) spanning typical regions involved in visual processing: left calcarine gyrus, cuneus, lingual gyrus and occipital superior and middle gyrus. These gamma power changes in visual areas did not vary with material (interaction p=0.42, t(19) =-0.81).

In accordance with MEG results, a significant number of electrodes showed significant gamma power increases related to memory formation in iEEG data (FEA: p_corr_ =0.0001, 43 electrodes, Figure 4G). Contrary to the MEG results, a small but significant number of electrodes showed a decrease in gamma power for later remembered items (FEA: p_corr_ =0.036, 27 electrodes, Figure 4G). This negative gamma SME could have been missed in MEG because of the limited spatial resolution and overall slightly worse signal to noise ratio in high frequencies, or could be a false positive in the iEEG data due to the inflated statistical power of the fixed effect analysis. Concerning the material specificity of memory effects however similar results as in MEG data were again observed: Gamma power SMEs did not vary with material (interaction material x memory p=0.39, t(9)=0.91, 10 patients with 2-13 electrodes, Figure 4F). Similarly, no significant influence of material on SMEs was found in the fixed effects analysis for the gamma frequency range (FEA: SME words < SME faces: p_corr_ =0.24, 21 electrodes, SME words > SME faces: p_cor_r =0.75, 13 electrodes, Figure 4G).

Results from both iEEG and MEG showed that increases in gamma power index successful memory formation independently of material, thus mirroring the results of theta power. To conclude, fixed effects and random effects analysis of iEEG data fully replicates the findings obtained in MEG: negative alpha/beta SMEs vary with material, positive gamma SMEs and negative theta SMEs are found irrespective of material. This frequency specific pattern of SME material dependency demonstrates that changes in specific frequency bands index specific processing demands during encoding.

### Latency differences between Gamma and Theta SMEs

The analyses described above demonstrate a functional dissociation of decreases in low frequency and increases in higher frequency power, suggesting that these changes do not reflect a single process. Specifically, alpha/beta power decreases varied with material during memory formation, whereas theta power decreases and gamma power increases did not. However, power decreases in theta and power increases in gamma did show a similar pattern of SMEs, i.e. both accompany memory formation independent of material. Therefore, in a further analysis we focused on the relationship between these two frequency bands. More specifically, we asked of whether theta power decreases and gamma power increases occur in the same regions and at the same time. In iEEG data, we calculated gamma SMEs in electrodes exhibiting significant negative theta SMEs and vice versa. Power spectra from all selected electrodes were averaged in each subject and entered into a dependent t-test (with N subjects as random variable). In electrodes exhibiting a significant positive gamma SME, no significant negative theta SME was evident in the early time window, in which gamma power changes were observed (Figure 5A, B, 2-5 Hz, 0.3 -1.0 sec, t(9)=-1.162, p=0.275). However, in a later time window, theta power significantly decreased for remembered vs forgotten items (2-5 Hz, 1.0-1.5 sec, t(9)=-3.137, p=0.0120, Figure 5B). This result indicates that theta and gamma SMEs can be located at the same regions but are shifted in time (i.e. gamma SME precedes theta SME).

**Figure 5.**
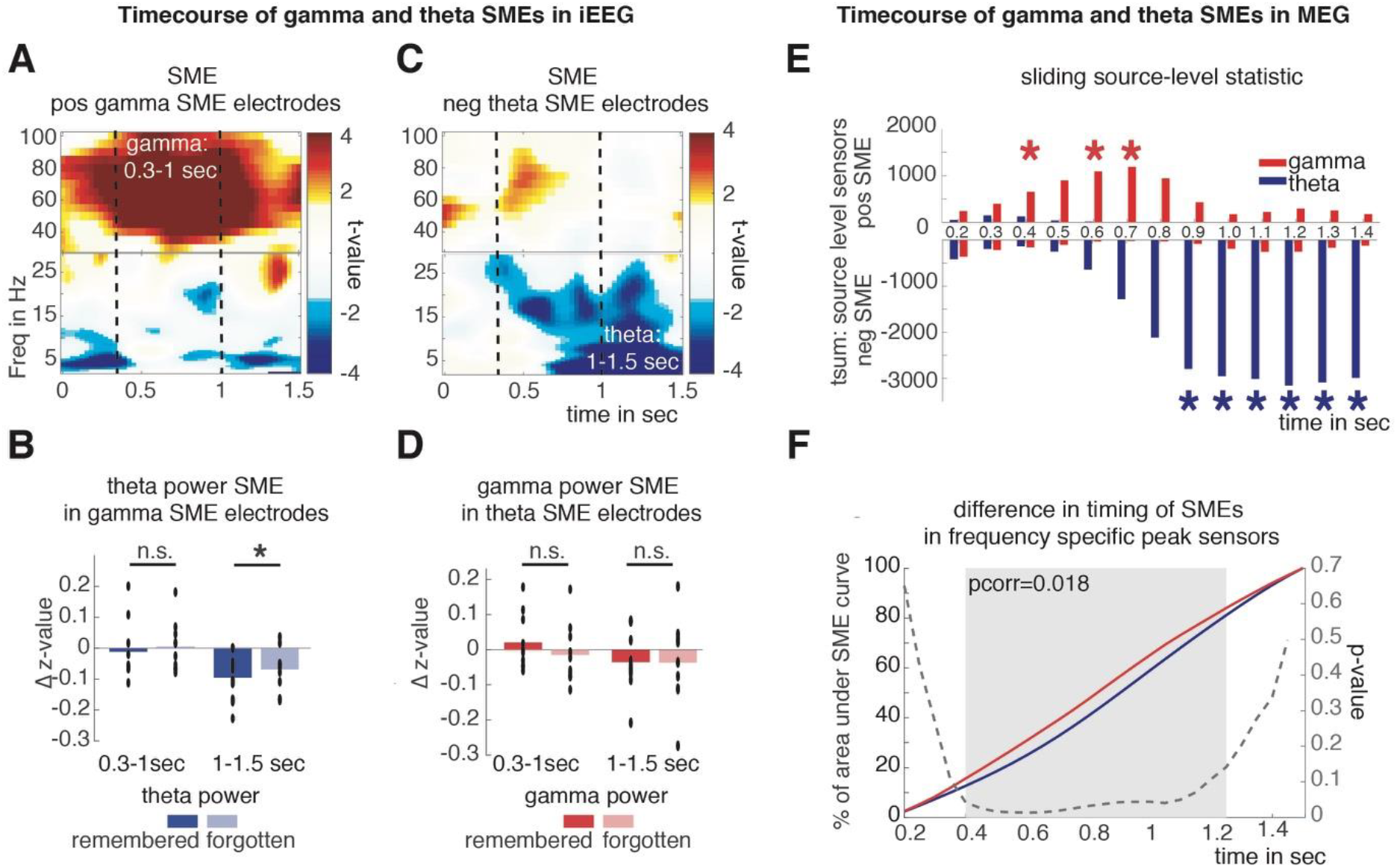
Timing differences of theta and gamma SMEs. (A –D) show differences in timing in iEEG data. Time-frequency plots depict t-values of SMEs in iEEG electrodes exhibiting a significant positive SME in the gamma frequency range or a significant negative SME in the theta frequency range respectively. Dashed lines highlight the time windows of gamma SMEs (0.3-1 sec) and theta SMEs (1.0-1.5 sec). (B) Bar plots show average theta SMEs in electrodes defined by gamma SMEs and gamma SMEs in electrodes showing theta SMEs for the earlier gamma and later theta time window. Black dots show the respective single subject averages, black asterisk mark significant differences (p<0.05 (E, F) depict differences in timing of theta and gamma SMEs in MEG. (E) Bars show the t-sum across significant negative and positive source level sensors of gamma and theta SMEs calculated on power averages in a 300 ms windows with 100 ms increments. Red and blue asterisks mark significant positive gamma SMEs of negative theta SMEs at the given timepoint respectively. (F) For each MEG dataset the 5% source level sensors were identified that exhibited the highest frequency specific SMEs across 0.2 to 1.5 sec. The average area under the SME curve reached at each timepoint in these frequency and subject specific ROIs is plotted for gamma and theta SMEs, the grey area highlights the significant cluster (random cluster permutation) in which gamma SMEs significantly precede theta SMEs.

This same analysis was repeated, now investigating whether gamma SMEs can be found in electrodes exhibiting significant theta SMEs (Figure 5C). Again, no significant gamma SMEs were evident at the same time window when theta SMEs were predominant (50-90 Hz, 1.0-1.5 sec: t(9)=0.058, p=0.958, Figure 5D). However, in the early time window there was a trend for a significant gamma SME, which did not reach significance (50-90 Hz, 0.3-1.0 sec, t(9)=1.785, p=0.11, Figure 5D). This result agrees with the preceding analysis in suggesting that theta and gamma SMEs may overlap spatially but can be temporally dissociated.

Intracranial EEG data are inherently restricted to the clinically defined implantation sites, and therefore one cannot exclude that theta and gamma effects overlap in areas where no electrodes were implanted. This limitation does not exist for whole brain recording methods like MEG. Therefore, we investigated whether the general timing of theta and gamma SMEs on a whole brain level followed our previous findings. To provide a more time resolved depiction of SMEs, a cluster permutation statistic on the source data was calculated in sliding time-windows of 300 ms (with increments of 100 ms) in the gamma frequency band (50-90 Hz) and the theta band (2-5 Hz). In Figure 5E the respective positive and negative t-sums across all significant source sensors are plotted. From this plot it is evident that positive gamma SMEs appear earlier than negative theta SMEs. To formally quantify these latency differences between gamma SMEs and theta SMEs, we calculated the percentage of the area-under-the-curve at each timepoint by cumulating the SME curve across time for each subject and at frequency specific peak source sensors (5% absolute highest gamma or theta SME summed across 0.2 to 1.5 sec). This analysis allows for an unbiased contrast of latency between SMEs of the two frequencies. Each frequency starts at the same point (0%) and finishes at the same point (100%). Analogously, this analysis is like pitching two horses against each other in a race and measuring the distance covered at each time point to see which one is faster. This area-under-the-curve method is more sensitive to detect differences in timing than classical peak latency comparisons, and is not affected by differences in time-resolution between gamma and theta SMEs. In Figure 5F the area-under-the-curve for negative theta SMEs and positive gamma SMEs is shown for each time bin from 0.2-1.5 sec. The latency curve for positive gamma SMEs is leading in respect to the theta SME curve indicating gamma SMEs earlier in time compared to theta SMEs. A cluster permutation statistic permuting the individual latency curve across subjects showed a significant difference in the timing of gamma and theta SMEs (cluster permutation p_corr_=0.018).

Together, these results illustrate that low frequency negative SMEs can be co-located to similar regions where positive SMEs in higher frequencies are observed. However, the SMEs in the gamma and theta frequency bands are temporally dissociated, with increased gamma power preceding decreases in theta power during the encoding of subsequently remembered items. This latter result indicates that the two frequency bands index different cognitive processes that occur at different time points in the service of memory.

## Discussion

The present results support the hypothesis that power changes in different frequency bands during memory encoding represent distinct fingerprints of differentiable encoding processes. In line with prior studies, decreases in theta and alpha/beta power and increases in gamma power predicted subsequent memory. Unlike the reported uniform decreases in low frequency power and increases in high frequency power predicting memory formation [8,20], the present results demonstrate that theta, alpha/beta and gamma power changes during encoding exhibit dissociable characteristics inconsistent with the frequency tilt hypothesis: (i) Areas involved in word and face processing exhibit material dependent decreases in alpha/beta power, while concurrent increases in gamma power were evident only in posterior sensory processing regions but not in left lateralized areas responsive to word processing, (ii) Negative alpha/beta SMEs depend on material and are stronger for words than faces, while negative theta SMEs and positive gamma SMEs do not depend on material, (iii) Negative theta SMEs and positive gamma SMEs differ in timing with gamma SMEs preceding theta SMEs. A particular strength of this study is that we show the same pattern of results in two independent datasets recorded in MEG and in iEEG. The reported dissociations are therefore present at different spatial scales and in independent of how the signal is being measured.

Our results replicate the commonly found pattern of low frequency power decreases and high frequency power increases during memory encoding. However, in-depth analysis of this pattern revealed functionally, spatially and temporally distinct signatures exhibited by different frequency bands. A simple analysis of time averaged power spectral density [11,17] as proposed for analyzing spectral tilts in the data would have concealed the diverse and frequency- and time specific patterns underlying memory formation. That said, summarizing complex changes in the power-spectrum in simpler metrics like tilt or broadband shifts clearly offers useful tools for characterizing overall brain states [47,48] especially regarding pathological states or age-related changes [11]. However, these metrics have a risk of concealing more fine-grained temporal and spectral dynamics underlying complex cognitive processing, like memory formation. To understand the neural dynamics involved in cognition, the explanatory value of spectral tilts or broadband shifts remains therefore limited. The specific relation of alpha/beta deceases to the encoding of words and the differences in timing of theta and gamma SMEs demonstrate that decreases in lower frequencies and increases in higher frequencies are not a uniform, ubiquitous marker of memory encoding, but instead index separable processes, which likely have explanatory value to understand transformation of experiences in durable memory traces.

Alpha/beta power in both iEEG and MEG specifically varied depending on the type of material to be encoded. Alpha/beta SMEs were stronger for the encoding of words in contrast to faces, i.e. material differing with respect to semantic processing properties. This interaction is specific to the alpha/beta band and thus cannot be explained from a spectral tilt perspective. If successful encoding of words is simply related to a “flatter” tilt of the power spectrum we should also see similar SMEs specific for words in the theta band, and mirroring SMEs in higher frequencies. However, in both datasets, MEG and iEEG, theta SMEs did not show the same interaction pattern as alpha/beta SMEs.

The present findings are consistent with frameworks that propose specific roles of different frequency bands i.e. spectral fingerprints of cognitive processing [26,27] of different frequency bands to multiplex content specific memory processes [49]. The dissociable roles of gamma and alpha/beta band oscillations have been studied in attention tasks [12,50]. Gamma oscillations in general have been hypothesized to play an important role in local bottom up sensory processing, whereas changes in alpha/beta oscillations have been related to long range cortical communication and top down processing [12,51,52]. This view of bottom up and top down processes are also in line with the current findings: gamma power increases were confined to occipital sensory areas and evident early after the stimulus, whereas decreases in alpha/beta and theta power were evident in widespread distributed cortical and medio-temporal regions in a later time window reflecting possibly higher-level processes involving top down control. The pattern of distinct spectral fingerprints in the present data concurs with cognitive memory models like Tulving’s SPI model [29] or PIMMS [30], which hypothesize memory encoding not as one monolithic process but highlight the role of different subprocesses. Our results show dissociable spectral fingerprints, which resemble commonly assumed stages in these models: perceptual, semantic and episodic processes.

Previous behavioral research has shown that words are especially well remembered if semantically processed [39], whereas memory for unfamiliar faces does not benefit from semantic encoding strategies [40]. We demonstrate that decreases in alpha/beta power specifically index encoding of verbalizable material. This finding agrees with previous studies showing that decreases in alpha/beta power are specifically related to semantic encoding [36,37,53]. Beta oscillations in particular have been connected to language and semantic processes [54,55]. On a more general level alpha/beta decreases have been linked to cortical information processing [56], or to distributed top down networks [57,58]. This view is also in line with prior studies that reported complex item specific representations being coded in the alpha/beta frequency range [59–61]. Because of these hypothesized features, alpha/beta oscillations might be a specific marker of the neural mechanisms behind the processing of distributed semantic features [62]. In the light of the present results and prior studies alpha/beta decreases during memory formation seem to be specific spectral fingerprints of semantic processing. The higher memory performance for words compared to faces on a cognitive level also highlights the important role of the semantic system as the main route to episodic memory [63]. The specific relationship of alpha/beta power decreases to the encoding of verbalizable material is a promising first step to a deeper understanding of the interaction of semantic and episodic memory.

Decreases in theta power and increases in gamma power related to subsequent memory are unspecific for the material that is to be encoded. Theta power decreases were spanning widespread cortical areas including MTL, frontal lobe, temporal, and parietal areas resembling the core memory network [64] and matching prior results [8,34]. Changes in the gamma band during encoding were specifically located in cortical areas involved in sensory visual processing, in line with previously reported SMEs in visual cortex [65] and in ventral occipito-temporal regions [8,23]. The insensitivity of the theta and gamma power changes to material and the general prominence of theta power changes in memory encoding studies suggest a role of theta decreases as a marker of MTL related memory encoding mechanisms independent of how stimuli are processed.

A careful analysis of the temporal dynamics of the gamma and theta SMEs revealed dissociable time courses of these effects. SMEs in gamma power preceded SMEs in theta (Figure 5). Such a latency difference is in line with the view of gamma power indexing early perceptual processing stages whereas theta power changes reflect later MTL related memory processes. Theoretical models of memory put these two computations usually at opposing ends of the processing cascade [29,30]. Methods that average power spectra across time are unable to detect these differences [17,20] and lose a major advantage of electrophysiological recordings, i.e. its high temporal resolution. Together, the timing difference between gamma and theta SMEs demonstrates that theta power decreases and gamma power increases reflect different neural processes which occur in sequence during memory encoding.

Our findings show that low frequency decreases and high frequency increases do not necessarily overlap in time and space. In addition to the differences in timing between gamma and theta SMEs, gamma power increases did not consistently occur in the same brain regions as alpha/beta power decreases during processing of words. While gamma power increases co-occurred with alpha/beta power decreases in occipital/posterior regions [66], no such co-localization was evident in left lateralized areas involved in semantic processing. Consequently, if the analysis in our study was limited to increases in broad band gamma power changes, a common practice in iEEG analysis [21–25][9,21,22] the prominent changes in alpha/beta power related to word processing would have been missed. Our findings are exemplary in demonstrating that limiting analysis to very narrow parts of the time-frequency spectrum restricts the potential insights that can be drawn from the data.

A unique aspect of the present study is that oscillatory brain activity was measured with two modalities, and in two separate subject groups. Intracranial recordings in patients with epilepsy and MEG recordings in a healthy student population exhibited concordant pattern of results. Importantly, both MEG and iEEG come with their own specific strengths and limitations [42,43]. MEG sensors, for instance, have different noise levels as a consequence of the sensors being mounted in a dewer, which results in different distances of sensors from brain tissue and different susceptibility to movement artefacts. These problems could affect signal-to-noise ratio at certain brain areas and frequency ranges, making it for example difficult to detect gamma effects in frontal regions. This problem does not exist with iEEG where electrodes are implanted directly in the brain tissue, thus allowing one to record electrophysiological activity in all frequency bands with high spatial resolution. MEG has two major advantages over iEEG: (i) whole head coverage in all subjects, and (ii) activity is recorded in healthy participants rather than a patient population. Although often seen as the gold-standard for electrophysiological studies in humans, iEEG data inherently suffers from the fact that the data is recorded in a non-healthy brain, possibly confounded with changes in frequency spectra, epileptogenic changes and artifacts [67,68]. Here we sought to combine the two methods in a complementary manner in order to overcome their respective limitations. Indeed, the tight overlap between iEEG and MEG results described here lends confidence in both results, in that the iEEG data verifies the MEG source reconstruction, and the MEG data alleviates concerns about the generalizability of the iEEG findings. Prior studies have combined M/EEG and iEEG, albeit typically in very small samples [32,33]. The present study is, to the best of our knowledge, the first to combine MEG and iEEG data of comparable sample sizes to study oscillatory processes in memory encoding. The present results therefore offer a unique validation of the commonly implied interchangeability of MEG and iEEG results.

An open question that remains is why decreases in theta power are involved in memory encoding in general, while alpha/beta power decreases are specifically involved in the encoding of semantically meaningful material. Arguably, both processes involve the integration of representations coded in widespread neural networks. Interestingly, power decreases which indicate a decrease in local synchronization are often found to co-occur with increases in long range phase synchronization [8, 69–71]. Therefore, decreases in power (i.e. local connectivity) might be a prerequisite for the formation of largescale fine-grained connectivity vital for distributed neural representations. The present findings thereby open up important follow-up questions concerning the relationship of local power decreases and network connectivity.

To conclude, we recorded electrophysiological activity during memory encoding in two complementary modalities, MEG and intracranial EEG. Our data provide evidence that the reported decreases in low frequency power and concurrent increases in high frequency power during memory encoding are not general markers of neural activity. Instead, the different responses in low and high frequency bands reflect spectral fingerprints of dissociable memory encoding processes. Considering interactions of different memory system networks, the specific relationship of alpha/beta power changes to semantic processing opens a window to the relationship of semantic and episodic memory, that has, as yet, not been well studied. Speculatively, interactions of alpha/beta and theta networks might specifically mark the interplay between semantic and episodic memory, which crucially shapes human memories [29,30].

## Material and Methods

### Participants

Thirty-two volunteers participated in the MEG experiment (compensated with course credit or 10 €/hour). Data from eleven participants was excluded because of low trial numbers in one of the conditions (minimum 30 trials) after rejecting MEG artifacts and trials with early response button presses. One additional dataset was excluded because of an erroneous headshape digitization, resulting in a sample of twenty subjects (M=23.5 years, range 18-33 years, 6 male). All subjects were right handed, spoke German as their native language, reported no history of neurologic or psychiatric disease, and had normal or corrected to normal vision.

Additional twenty-two patients with pharmaco-resistant epilepsy implanted with intracranial electrodes for diagnostic purposes volunteered to participate in a matching memory encoding study. Data of seventeen patients were recorded at the University Hospital Erlangen, data of five patients were recorded in cooperation with University College London at the National Hospital for Neurology & Neurosurgery. Data of eight patients were excluded from later analyses, due to either technical problems during recordings, left-handedness or insufficient memory performance. In the remaining dataset of thirteen patients (M=35.54 years, range: 20-60 years, 3 male) eight patients were native German speakers, four native English speakers, and one Slovenian. Word material and instructions were translated accordingly.

All participants gave their written informed consent, and the experimental protocol was approved by the local ethical review board.

### Material

Word material was drawn from the MRC Psycholinguistic Database[72], translated into German/English/Slovenian depending on the respective participant (264 words during experiment, additional 12 words for practice trials). Neutral unfamiliar faces (264 faces during experiment, additional 12 faces for practice trials), were drawn from several face material databases ([73], and pics.stir.ac.uk). All face stimuli had an emotionally neutral expression and were presented in greyscale on a black background. Use of word and face material during encoding and as new material during recognition was counterbalanced across participants.

### Procedure

Every participant completed two task blocks: one block of word encoding and recognition and one block of encoding and recognizing unfamiliar faces (order counterbalanced across participants). During the whole experiment MEG or iEEG was recorded.

During the encoding phase participants were instructed to judge each item presented for pleasantness on a 1-6 scale. The encoding phase was followed by a distractor task to prevent working memory contributions to the recognition test. The distractor task was a variation of the inattentional blindness task as used in [74]. During the recognition phase all previously shown items were presented randomly intermixed with new items, i.e. lures. Participants were instructed to provide confidence ratings ranging from 1, very sure old, to 6, very sure new. Prior to each phase participants completed a short practice phase to familiarize them with the paradigm. In the MEG sample responses were given using two response boxes with 3 buttons placed on the right and left side of the body, response hand use was counterbalanced across participants. Due to the test setting in the hospital bed in the iEEG sample response hand use was not controlled, as flexible response hand use was not always possible. Patients completed the same paradigm as healthy controls with small adaptions (a reduced number of trials, self-paced responses and additional breaks, see Figure 1A).

### Behavioral analysis

A ROC approach was used to analyze memory performance. A single process unequal variance model was fitted to the data to obtain bias free measures of memory strength [75,76] and classify hits and misses relative to individually defined neutral response criteria for MEG/iEEG analysis (for details of fitting procedures see [35,37]). In short, this approach assumes that memory strength can be modeled by separate normal distributions for new and old items. The distance d’ of the mean of these distributions yields a bias free measure of memory strength. The model assumes that subjects respond with a certain confidence rating i, whenever their subjective memory strength exceeds a certain criterion c_i_. The crossover of the distributions of new and old items represents the point of the neutral response criterion as this point represents the memory strength that has an equal probability to be elicited by new and old items. An item that during recognition received a confidence rating i was classified as a hit if the corresponding estimated criterion c_i_ was higher than the individually estimated neutral criterion in the recognition block, otherwise the trial was classified as a miss. As demonstrated previously this procedure enhances signal to noise ratio by considering individual differences in the use of confidence ratings for hit and miss trial definition [35].

### MEG recording and processing

MEG was recorded with a 148-channel whole-cortex magnetometer (MAGNES 2500 WH, 4D Neuroimaging, San Diego, USA) in a magnetically shielded room, while participants were in a supine position. Data were continuously recorded at a sampling rate of 678.17 Hz, and bandwidth of 0.1-200 Hz. The participants’ nasion, left and right ear canal, and head shape were digitized prior to each session with a Polhemus 3Space Fasttrack.

All analyses were carried out in MATLAB (The MathWorks, Natick, MA) using the fieldtrip toolbox (www.ru.nl/fcdonders/fieldtrip, [77]). Data from encoding phases were epoched in trials −1.5 to 3 sec around each item onset during encoding. Line noise was removed by a discrete Fourier transform filter. Idiographic artifacts (channel jumps, muscle artifacts, noisy channels) were excluded from further analysis by visual inspection. Infomax independent component analysis was applied to correct for residual artifacts (e.g. eyeblinks, eye movements, heart beat related activity or tonic muscle activity). On average 105.8 word hit trials (SD=18.7, range: 75-129), 51.6 word miss trials (SD=18.4, range 31-88) and 85.8 face hit trials (SD=17.3, range: 39-111) and 71.8 faces miss trials (M=71.8, SD=18.0, range: 49-111) passed artifact correction. MEG sensor level data was transformed into planar gradients for sensor level analysis. This procedure emphasizes activity directly above a source, simplifying interpretation of MEG topographies [78]. Source analysis was carried out using a linearly constrained minimal variance (LCMV) beamformer [79], calculating a spatial filter based on the whole length of all trials. Individual structural MR images were aligned with the MEG sensor coordinates using NUT-MEG [80]. Individual single-shell headmodels [81] were constructed using structural MRIs of each participant. The brain space was divided in 10 mm grid voxels and normalized to the MNI brain using a warping procedure. Source time-courses for each grid point were calculated and subjected to a wavelet analysis described below.

Data was filtered to obtain lower frequency oscillatory power between 2 Hz and 30 Hz using wavelets with a 5-cycle length, resulting time-frequency data was smoothed with a Gaussian kernel (FWHM 200ms and 2 Hz) to account for inter-individual differences and changes in time-frequency resolution across frequencies. To obtain higher frequency oscillatory power in the gamma range (30-100 Hz) a multi-taper approach was used with a 300 ms window and a spectral smoothing of 10 Hz resulting in the use of 5 tapers. Resulting data was z-transformed to the respective mean and standard deviation across time for every time-frequency bin separately for two different recording blocks of words and faces but not separately for hits and misses (e.g. [8]).

### iEEG recording and processing

Intracranial data was recorded from subdural grid, strip, and depth electrodes (AdTech, recording system Deltamed, Natus or Nicolet, NicVue) referenced to a scalp electrode. The implantation scheme depended on the suspected epileptic foci and was therefore highly variable across patients (see Figure 1C). Locations of electrodes were determined using co-registered post-implantation MRIs and post-implantation CTs. Locations were then transformed to MNI coordinates by normalizing the post-implantation MRIs to standard MNI space using SPM8. Data were continuously recorded at different sampling rates (4 datasets: 512 Hz, 8 datasets:1024Hz, 1 dataset: 4096 Hz).

Data from encoding phases were epoched in trials from −1.5 to 3 secs around each item onset during encoding and downsampled to a sampling rate of 500 Hz. Data was referenced to bipolar montages to obtain maximally focal spatial resolution [82]. To this end, each electrode was re-referenced to its neighboring electrode (for grid electrodes across the horizontal and vertical dimension). Coordinates of these bipolar “virtual” electrodes were calculated as located between the respective physical electrodes. Electrodes within or bordering areas later resected or identified as the epileptic foci were excluded from analysis. Data was carefully visually inspected by a trained neurologist and a second individual; electrodes with epileptogenic activity were excluded from analysis. Individual trials which exceed the mean range, variance or kurtosis by more than 5 standard deviations were automatically rejected. Channels yielding less than 10 trials after artefact correction in any condition were rejected. This resulted in a dataset of 657 bipolar channels (of 926 recorded channels, M=51.85 per patient, std= 24.37, range 9-88). The mean number of trials across channels and patients for word hits was M=63.61 (std: 10.44 range: 47.16-83.72), for face hits M=47.84 (std 13.45 range: 19.15-68.70), for word misses M=27.39 (std 10.40 range: 12.61-43.45), and for face misses M=41.63 (std 15.56 range: 24.47-72). iEEG data was filtered to obtain oscillatory power, z-transformed, and smoothed using the same settings as MEG data.

### Statistical analysis: Memory and material effects

The study follows a 2×2 design with the factors Memory (remembered vs forgotten) and material (face vs word). The analysis scheme of iEEG and MEG data differed as a result of the different nature of these two datasets.

MEG data was analyzed using a conventional repeated measurements random effects design. Task contrasts of interest were interaction effects and main effects of the 2×2 repeated-measurements design (i.e., power spectrum for material x memory). In order to stay within the fieldtrip cluster statistic framework, these contrasts were calculated using the cluster permutated t-contrasts. In each subject power-spectra at each sensor/ source level sensor were averaged across all trials for each cell of the 2×2 design matrix (word-hits, word-miss, face-hit, face-miss). Main effects were analyzed by contrasting the means across the respective cells using dependent t-contrasts (i.e. mean of word-hit and words-miss vs mean of face-hit and face-miss as main effect of material). Interaction effects were calculated by contrasting the material specific SMEs (i.e. word-hit minus word-miss contrasted to face-hit minus face-miss). Averaging of cell specific means prevents a potential biasing of main effects by trial number differences across condition (i.e. more word-hit trials than face-hit trials. This analysis scheme of t-tests for testing interaction and main effects is equivalent to a 2 × 2 repeated-measures ANOVA.

Statistical analysis of MEG data was carried out using the fieldtrip cluster permutation approach[83]. The cluster permutation test consists of two steps: first, clusters of coherent t-values exceeding a certain threshold along selected dimensions (time, frequency, electrodes/grid voxels) are detected in the data. Second, summed t-values of these clusters are compared to a null distribution of t-sums of random clusters obtained by permuting condition labels across subjects. This procedure effectively controls for type I errors due to multiple testing. For sensor analysis three dimensional clusters (electrodes x time x frequency) were built by identifying neighboring time-frequency-channel bins involving at least two neighboring channels with a p-value below 0.01 (lower p-threshold to identify coherent clusters in higher dimensional clustering). For source space analysis clusters were formed across the spatial dimension (p-level threshold 0.05).

iEEG electrodes across the whole patient sample covered widespread brain areas, however the varying electrode implantation scheme in each patient impeded a similar random effects analysis as in MEG. iEEG analysis was restricted to the two time-frequency windows identified in MEG analysis (alpha/beta 8-20 Hz, 0.3-1.5 sec, gamma: 50-90 Hz, 0.3-1 sec) or a-priori defined (theta 2-5Hz, 1-1.5sec). A 2×2 ANOVA (memory x material) was calculated on the single trials in each single electrode. To evaluate the concordance of iEEG and MEG results, the overlap of iEEG and MEG effects was estimated by counting the number of iEEG electrodes showing a negative, positive or non-significant difference in each significant MEG source cluster. These absolute numbers of electrodes in each MEG defined region was used to construct a contingency table and a chi square test of independence was employed to assess statistical significance of the dependency of MEG and iEEG effects.

To elucidate differences in SMEs (i.e. interaction of material x memory) in iEEG data in a group random effects manner, all electrodes showing a significant main effect of memory in a frequency band of interest were averaged in each subject and subjected to the same random effects repeated measurement ANOVA analysis schema as MEG data. This random effects group analysis of an interaction was restricted to preselected electrodes exhibiting a main effect of memory. This preselection is no circular analysis (no “double dipping”), as interaction effects are independent of main effects.

To further ensure that the preselection of memory effect electrodes is not missing effects, an additional fixed effects analysis combining all electrodes across all patients was carried out. To this end a random distribution of the number of electrodes showing a negative or positive main or interaction across the whole patient/electrode sample was estimated by randomly shuffling the trial labels (word-face, remembered-forgotten) in each subject 10000 times and calculating the number of significant electrodes in each permutation. If the number of electrodes exhibiting a significant main or interaction effect in the data was observed in less than 5% of permutation, the pattern of results was regarded significant.

### Analysis of timing differences

For analysis of timing differences of gamma and theta SMEs in MEG we employed an area-under-the-curve-analysis, which has been shown to be more reliable in finding latency differences compared to peak latency analysis [84]. First, a t-contrast was calculated for theta and gamma power for all remembered vs forgotten trials in each subject for each source level sensor at each time-bin between 200-1500ms. Second, these t-value timecourses were thresholded at zero, for gamma SMEs all negative t-values were set to zero (as timing of positive SMEs was of interest) and vice versa for theta SMEs (all positive t-values were set to zero). Third, frequency specific peaks were identified by summing these thresholded t-value time-series across the whole time series and the selecting the 5% of sensors with the highest absolute t-sum. Fourth, the t-value time-series was averaged across the peak sensors in each subject and the area under the SME curve was calculated by integrating across the time dimension (i.e. summing at each timepoint the t-values up to this timepoint relative to the t sum across all time-bins). This procedure returns a time-resolved curve illustrating for each timepoint the percentage of the SME curve covered. To assess statistical differences in area-under-the-curve measure of theta and gamma SMEs, again a cluster permutation approach was utilized, shuffling the condition (theta/gamma) 1000 times across subjects and clustering significant t-values along the time dimension.

## Acknowledgments

We thank Sarang Dalal for help concerning the MEG source analysis and Gerd T. Waldhauser and Benjamin Griffiths for valuable discussion on the manuscript.

